# Experimental and stochastic models of melanoma T-cell therapy define impact of subclone fitness on selection of antigen loss variants

**DOI:** 10.1101/860023

**Authors:** Nicole Glodde, Anna Kraut, Debby van den Boorn-Konijnenberg, Saskia Vadder, Florian Kreten, Jonathan L. Schmid-Burgk, Pia Aymans, Kai Echelmeyer, Martin Rumpf, Jennifer Landsberg, Tobias Bald, Thomas Tüting, Anton Bovier, Michael Hölzel

**Affiliations:** Institute of Experimental Oncology, University Hospital Bonn, University of Bonn, Bonn, Germany; Department of Dermatology, University Hospital, Magdeburg University, Germany; Laboratory of Experimental Dermatology, Department of Dermatology, University Hospital Bonn, Bonn, Germany; Institute for Applied Mathematics, Bonn University, Bonn, Germany; Broad Institute of MIT and Harvard, Cambridge, MA 02142, USA; Department of Dermatology and Allergy, University Hospital Bonn, Bonn, Germany; Institute for Numerical Simulation, University of Bonn, Bonn, Germany; Oncology and Cellular Immunology Laboratory, QIMR Berghofer Medical Research Institute, Herston, Queensland, Australia

**Keywords:** Melanoma, immunotherapy, resistance, antigen loss, selection, clonal evolution, phenotypic plasticity, intratumor heterogeneity, Markov process, mathematical model, stochastic

## Abstract

Antigen loss is a key mechanism how tumor cells escape from T-cell immunotherapy. Using a mouse model of melanoma we directly compared antigen downregulation by phenotypic adaptation with genetically hardwired antigen loss. Unexpectedly, genetic ablation of Pmel, the melanocyte differentiation antigen targeted by adoptively transferred CD8^+^ T-cells, impaired melanoma cell growth in untreated tumors due to competitive pressure exerted by the bulk wild-type population. This established an evolutionary scenario, where T-cell immunotherapy imposed a dynamic fitness switch on wild-type melanoma cells and antigen loss variants, which resulted in highly variable enrichment of the latter in recurrent tumors. Stochastic simulations by an individual-based continuous-time Markov process suggested variable fitness of subclones within the antigen loss variant population as the most likely cause, which was validated experimentally. In summary, we provide a framework to better understand how subclone heterogeneity in tumors influences immune selection of genetic antigen loss variants through stochastic events.

## Introduction

Cytotoxic CD8^+^ T cells play an important role in tumor immune surveillance. They recognize peptides (epitopes) derived from tumor cell-encoded gene products (antigens) which are presented on major histocompatibility complex (MHC) class I molecules. Epitope-specific activation of CD8^+^ T cells leads to tumor cell killing through the release of cytotoxic granules and cytokines, amongst others, which can be exploited therapeutically by different strategies. One approach is genetic engineering of autologous CD8^+^ T cells by introducing a tumor antigen-specific T cell receptor (TCR) (D’Ippolito et al., 2019; Yang and Rosenberg, 2016). After *ex vivo* expansion these TCR-transgenic (TCRtg) CD8^+^ T cells can be re-infused into the same patient, a procedure known as adoptive T cell transfer (ACT). Clinical trials have been conducted in patients with various types of cancer showing that the emergence of resistant tumor cell variants restrains durable responses (Chodon et al., 2014; Mehta et al., 2018; Tran et al., 2016).

Malignant melanoma, an aggressive type of skin cancer, is a paradigm disease for the development of novel immunotherapies including ACT. Using mouse models, we previously found that melanomas can escape from ACT targeting the melanocyte differentiation antigen Pmel (also known as gp100) by dedifferentiation (Landsberg et al., 2012), an epigenetic mechanism also known as phenotype switching or phenotypic plasticity (Hoek and Goding, 2010; Hoek et al., 2008). Briefly, infiltrating activated Pmel-specific CD8^+^ T-cells (Pmel-1 T-cells) instigated pro-inflammatory cytokine release which induced dedifferentiation and downregulation of the Pmel antigen in melanoma cells, which impaired immune recognition and killing (Landsberg et al., 2012; Riesenberg et al., 2015). Reciprocally, melanoma cells upregulated mesenchymal and neural crest progenitor cell traits, a finding that was recently confirmed in melanoma patients treated with ACT directed against the differentiation antigen MART-1 (Mehta et al., 2018; Reinhardt et al., 2017). Thus, phenotypic plasticity of melanoma cells emerges as a relevant mechanism of resistance to immunotherapy (Arozarena and Wellbrock, 2019; Hölzel et al., 2013).

A better understanding how the various cell populations interact over time could help to improve current ACT regimens, but longitudinal analyses impose challenges with regard to tissue sampling, both in patients and animal models. We therefore reasoned that mathematical modeling as in (Eftimie et al., 2011; Kuznetsov et al., 1994) could complement our experimental approaches and potentially make novel predictions. Recently, we proposed a stochastic model of ACT, which is an extension of individual-based stochastic models for adaptive dynamics, where we use the term individual for cells and relevant events for each individual (e.g. cell division and death) occur randomly (Baar et al., 2016). Using adjusted parameters the model faithfully recapitulated tumor growth kinetics and melanoma cell state transitions as reported in our previous experimental study (Landsberg et al., 2012).

Here, we used experimental and mathematical models of ACT in order to compare the evolution of gene edited Pmel antigen loss variants (Pmel knockout; Pmel^KO^) with adaptive dedifferentiation, because genetically hardwired loss of target antigen expression or presentation is another key mechanism of resistance to immunotherapy (Restifo et al., 1996; Sucker et al., 2017; Zaretsky et al., 2016). ACT strongly selected for Pmel^KO^ melanoma cells, but unexpectedly they exhibited a growth defect in the absence of therapy imposed by the bulk wild-type (WT) melanoma cell population. This established an evolutionary scenario of competing tumor cell populations, where Pmel^KO^ and WT melanoma cells switch their fitness in response to ACT. Accounting for context-dependent fitness, the stochastic model suggested subclone fitness variability as the most likely cause for the unanticipated high variability of Pmel^KO^ cells enrichment found in ACT-recurrent melanomas, which was confirmed experimentally by different approaches. Our work emphasizes the need to take evolutionary dynamics into account when interpreting variant allele frequencies in therapy resistant tumor specimens and it provides a framework for stochastic modeling these scenarios.

## Results

Our experimental ACT regimen consists of (i) a single dose of cyclophosphamide (chemotherapeutic conditioning), (ii) intravenous injection of Pmel-specific CD90.1^+^CD8^+^ TCRtg Pmel-1 T cells, (iii) *in vivo* activation of Pmel-1 T cells with a human gp100-expressing adenovirus vaccine (Ad-hgp100) and (iv) innate immune system activation with CpG and poly(I:C) (Kohlmeyer et al., 2009; Landsberg et al., 2012). Recently, we found that short-term concomitant treatment with small molecule inhibitors (e.g. capmatinib) of the c-MET receptor tyrosine kinase (METi) enhanced ACT efficacy (ACT^METi^) by blocking the recruitment of T cell-suppressive neutrophils early during treatment, regardless whether c-MET signaling was an intrinsic driver of melanoma cell growth (HCmel12 model) or not (B16F1 model) (Glodde et al., 2017).

### Dynamic Pmel antigen expression in response to MET inhibitor (METi) treatment

The syngeneic HCmel12 melanoma model was established from a primary melanoma in *Hgf-Cdk4^R24C^* mice (Bald et al., 2014), a model in which transgenic expression of the ligand HGF (hepatocyte growth factor) constitutively activates c-MET signaling. As HCmel12 grows rapidly, phenotype switching occurs dynamically and genetic manipulation is facile (Bald et al., 2014), we considered HCmel12 as being suited for modeling the dynamics of different melanoma cell populations during ACT^METi^. We first asked whether METi treatment could influence the phenotype (differentiation) of HCmel12 cells, an important parameter in our stochastic ACT model (Baar et al., 2016), because it was reported that MAPK inhibitors upregulated melanocyte differentiation antigens in human melanomas (Boni et al., 2010; Frederick et al., 2013). In order to have matched METi-insensitive controls in addition to B16F1, we generated METi-resistant variants of HCmel12 (HCmel12^METi-R^), which remained METi-resistant even after passaging *in vivo* and showed sustained c-MET downstream signaling activity (phospho-ERK and phospho-AKT) in the presence of METi (Figure S1A-D). Indeed, we found that METi (capmatinib) induced Pmel protein expression in parental HCmel12 cells (WT, wild-type), whereas it remained unchanged in B16F1 cells and HCmel12^METi-R^ (Figure S1E). Also METi treatment impaired TNF-α induced dedifferentiation of HCmel12 WT cells, but not B16F1 cells (Figure S1F). Despite these dynamic changes of Pmel, concomitant METi treatment enhanced ACT efficacy against HCmel12^METi-R^ melanomas and Pmel-1 T-cell expansion to the same extent as previously reported for parental HCmel12 WT melanomas (Figure S1G-J) (Glodde et al., 2017). In conclusion, short-term METi treatment increased Pmel antigen levels in HCmel12 WT cells, but this effect did not account for the immunomodulatory effects of METi treatment in conjunction with ACT in line with our previous work. An explanation is that the vaccination with Ad-hgp100 transiently provides excess antigen to the adoptively transferred Pmel-1 T cells, which limits the influence of Pmel antigen level expressed by tumor cells during the early phase of treatment. However, we reasoned that antigen expression by tumor cells becomes a critical determinant later in the course of therapy when adenoviral antigen expression declines.

### Highly variable enrichment of Pmel^KO^ variants in ACT^METi^ recurrent melanomas

To confirm this, we analyzed ACT^METi^-recurrent HCmel12 melanomas and we found reduced expression of Pmel by immunohistochemistry (Figure 1A-C), in line with melanoma dedifferentiation being a mechanism of ACT resistance (Landsberg et al., 2012; Mehta et al., 2018). This in turn raised the question whether genetic ablation of *Pmel* in HCmel12 cells would confer any additional advantage to escape from ACT^METi^. Using CRISPR-Cas9 we targeted the *Pmel* gene upstream of the region encoding for the gp100 epitope with three different sgRNA constructs for control purposes, which were transfected individually into HCmel12 cells. Co-transfection of a GFP expression construct allowed us to isolate highly transfected cells in order to increase the frequency of HCmel12 Pmel^indel^ cells in polyclonal cell cultures. After a short period of *in vitro* expansion we determined the Pmel^indel^ allele frequencies by amplicon NGS using a modified version of the OutKnocker tool (Schmid-Burgk et al., 2014), which revealed polyclonal indel distributions typical for CRISPR-Cas9 gene editing with largely inactivating frame-shift events (Figure 1D).

**Figure 1.**
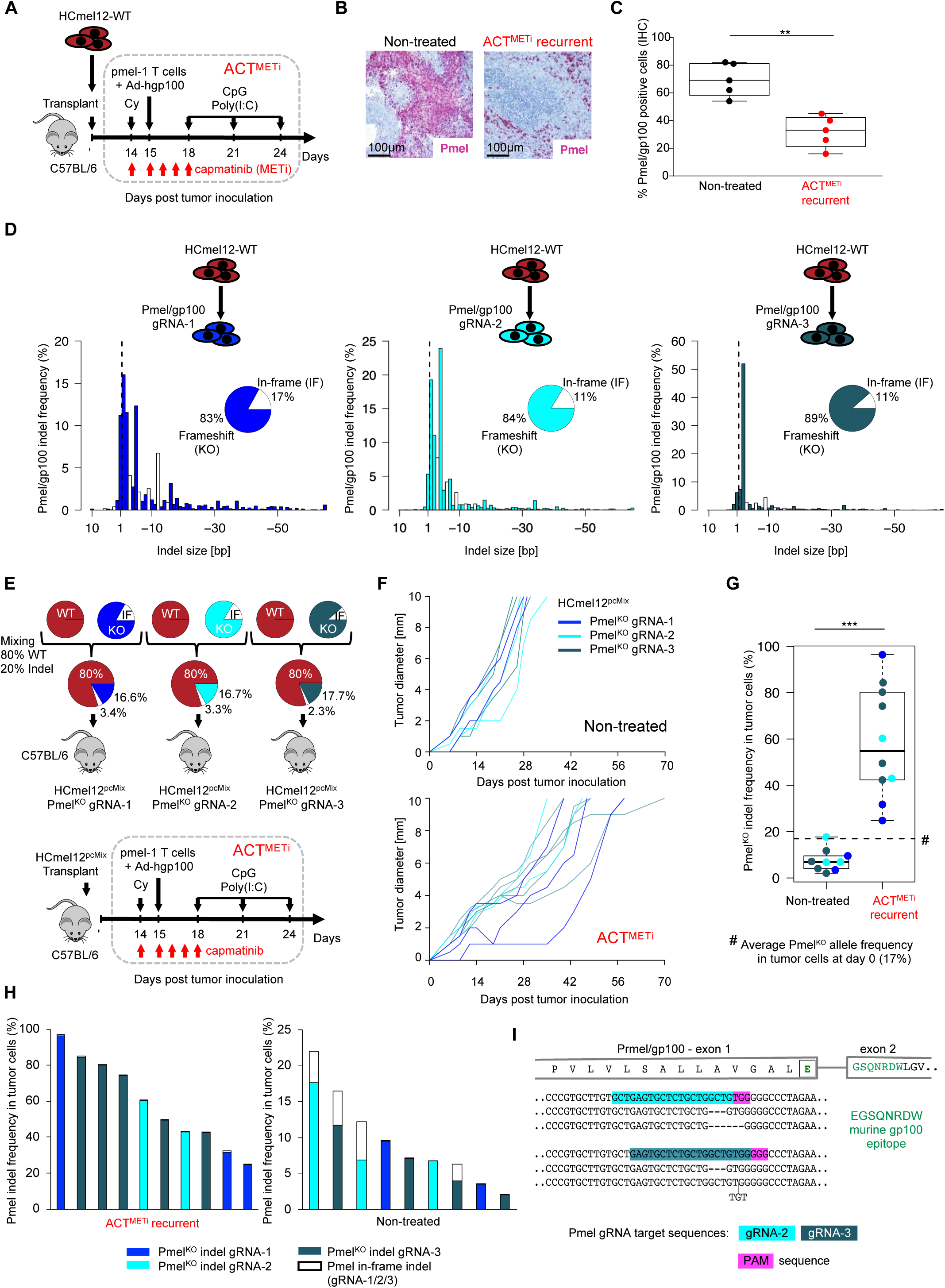
Highly variable enrichment of Pmel^KO^ antigen loss variants in ACT^METi^-recurrent melanomas. **(A)** Experimental outline of HCmel12 WT melanomas treated with ACT^METi^. **(B)** Representative Pmel expression by immunohistochemistry in untreated and ACT^METi^-recurrent HCmel12 WT melanomas. **(C)** Quantification of the experiments from B. Two-sided Mann-Whitney-U test. **(D)** Pmel indel distribution in HCmel12 cells gene-edited with the indicated sgRNAs. Portions of frameshift and in-frame indels are indicated. **(E)** Experimental outline of HCmel12^pcMix^ melanomas treated with ACT^METi^. **(F)** Individual tumor growth curves of HCmel12^pcMix^ melanomas left untreated or treated with ACT^METi^. **(G)** Amplicon NGS based quantification of Pmel^KO^ allele percentages in the tumor cell fraction of untreated or recurrent ACT^METi^-treated HCmel12^pcMix^ melanomas. **(H)** As G, but total Pmel indel percentages and the respective portions of frameshift and small in-frame indels. **(I)** Examples of small in-frame indels detected in G, H.

We then mixed the three polyclonal HCmel12 Pmel^indel^ cultures with parental HCmel12 cells (WT; wild type) cells in order to obtain a Pmel^indel^ allele frequency of 20% and injected these mixtures (HCmel12^pcMix^; polyclonal mix culture) into the flank of syngeneic mice (Figure 1E). Animals bearing established HCmel12^pcMix^ melanomas were treated with ACT^METi^ or left untreated for control purposes (Figure 1F). Of note, all HCmel12^pcMix^ melanomas escaped from ACT^METi^, whereas we previously reported that ACT^METi^ achieved eradication of parental HCmel12 melanomas in about 30% of cases (Glodde et al., 2017). When tumors reached a size of 10mm in diameter and mice had to be sacrificed for animal welfare, we isolated genomic DNA from untreated as well as recurrent HCmel12^pcMix^ melanomas and we determined the Pmel indel allele frequencies by amplicon NGS. A HCmel12-specific mutation in p53 (p53^R172H^) was used for tumor cell content normalization, as this mutation is absent from host cells in the tumor microenvironment. Pmel indel alleles were strongly enriched in recurrent HCmel12^pcMix^ melanomas being on average >99% inactivating (Pmel^KO^) frame-shift indels (Figure 1G), which demonstrated that HCmel12 Pmel^KO^ cells gained fitness in comparison to HCmel12 WT cells in response to ACT^METi^. Notably, this enrichment was subject to unanticipated high variability ranging from 20% to almost 100% (mean 58.5%).

### Reduced fitness of Pmel^KO^ variants within a bulk wild-type melanoma cell population

Furthermore, the average Pmel^KO^ allele frequency detected in untreated HCmel12^pcMix^ tumors (mean 7.8%) was clearly below the frequency at the time of injection (17%) (# in Figure 1G), which again was unexpected. This suggested that HCmel12 Pmel^KO^ cells had a reduced fitness in the context of a bulk HCmel12 WT population when immunological selection pressure by Pmel-1 T cells was absent. To corroborate this notion, we analyzed the dynamics of small in-frame indel alleles that are likely to preserve Pmel protein function. We hypothesized that HCmel12 cells with small in-frame indels should exhibit the same context-dependent fitness as HCmel12 WT cells. In support for this idea, several untreated HCmel12^pcMix^ melanomas contained a substantial proportion of small in-frame indel alleles (up to 44% of total Pmel indels), whereas small in-frame indel alleles were almost undetectable (<1% of total Pmel indels) in ACT^METi^-recurrent HCmel12^pcMix^ melanomas (Figure 1H and 1I). In conclusion, this finding supported the notion that genetic Pmel loss reduced HCmel12 fitness in untreated melanomas, but strongly increased their fitness under ACT^METi^.

### Wild-type melanoma cells impose competitive pressure on Pmel^KO^ melanoma cells

Next, we analyzed the *in vivo* growth rate of pure HCmel12 Pmel^KO^ melanomas, because a growth rate similar to HCmel12 WT melanomas would argue that HCmel12 WT cells exerted competitive pressure on Pmel^KO^ cells within HCmel12^pcMix^ melanomas. For this purpose, we isolated three independent HCmel12 Pmel^KO^ clones and validated Pmel gene editing by amplicon NGS as well as Western blot confirming absence of Pmel protein expression (Figure S2A and S2B). An equal mixture of the three HCmel12 Pmel^KO^ clones was injected into the flank of syngeneic mice and indeed pure HCmel12 Pmel^KO^ melanomas exhibited a growth rate similar as HCmel12 WT melanomas (Figure S2C). Hence, this result argues for a scenario where HCmel12 WT cells impose a competitive pressure on HCmel12 Pmel^KO^ cells.

### Stochastic model of ACT^METi^ implementing pre-existing Pmel^KO^ antigen loss variants

Our herein described approach with HCmel12^pcMix^ melanomas established an evolutionary scenario of competing tumor cell populations undergoing a reciprocal fitness switch upon ACT^METi^. This prompted us to use mathematical modeling in order to better understand of cell population dynamics and potentially explain the highly variable enrichment of Pmel^KO^ cells in recurrent melanomas. The stochastic model of ACT^METi^ is based on the individual-based Markov process in Baar et al. (Baar et al., 2016). We adjusted this model according to the context of HCmel12^pcMix^ melanomas treated with ACT^METi^. We consider six different types of interacting cell populations or molecules collectively termed as individuals: Differentiated (Diff) and dedifferentiated (Dedi) WT melanoma cells, Pmel^KO^ melanoma cells (KO), CD8^+^ Pmel-1 T-cells (CD8), cytokines (Cyto), and dead melanomas cells (Dead). The latter are included, because we reasoned that they contribute to measured tumor sizes, although they do not influence the evolution of the other cells in our mathematical model. Cytokines comprise a variety of different molecules, in particular T cell effector cytokines such as TNF-α and IFN-γ, that evoke a pro-inflammatory microenvironment promoting melanoma cell dedifferentiation and upregulation of negative immune checkpoint molecules (Chen et al., 2019; Landsberg et al., 2012; Reinhardt et al., 2017; Riesenberg et al., 2015). The state of the mathematical process

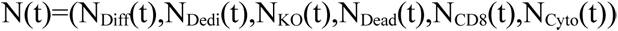

describes how many of these different types of individuals (cells, cytokines) are present in the tumor tissue as a function of time t starting with tumor cell inoculation at t=0.

We distinguish different events that change the state of this system: Cell division, cell death, cytokine secretion, killing of WT^Diff^ melanoma cells, phenotype switching between WT^Diff^ and WT^Dedi^ melanoma cells, and spontaneous mutations generating Pmel^KO^ melanoma cells. All of these events happen at certain rates or frequencies that depend on several fixed parameters and the current state of the system. The dynamics of the Markov process are summarized by its infinitesimal generator that takes the form of

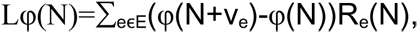

where E is the set of possible events, v_e_ is the change in the population associated to an event, and R_e_(N) is the rate at which the event occurs. It can be constructed rigorously similar to the process in (Fournier and Méléard, 2004). The following examples illustrate two events: (i) Increased cytokine release results in a higher rate of dedifferentiation of WT^Diff^ melanoma cells, in which case the cytokine induced switch happens at rate 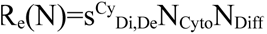; (ii) an increase in WT melanoma cells imposes a higher competitive pressure (parametrized by c_KO,Di_/c_KO,De_) on Pmel^KO^ cells elevating their rate of cell death R_e_(N)=(d_KO_+c_KO,Di_N_Di_+c_KO,De_N_De_+c_KO,KO_N_KO_)N_KO_. The effects of ACT^METi^ are modeled through (i) the addition of 10^5^ Pmel-1 T-cells on the respective day of adoptive T-cell transfer and (ii) the intermittent change of parameters within the five days of concomitant METi treatment, namely reduced WT melanoma cell dedifferentiation, a general reduction of tumor cell growth through a decreased cell division rate, and an increased rate of Pmel-1 T-cell proliferation and killing of WT^Diff^ cells via decreased T-cell suppression based on experimental data in Glodde et al. Figure 2A shows an interaction diagram that summarizes all these effects. A more detailed description of the mathematical model and the different events and rates is given in the methods section. Simulations using the adjusted model recapitulated the tumor growth kinetics of HCmel12 WT melanomas treated with ACT^METi^ from Glodde et al. (Figure 2B and 2C). Of note, our simulations also predicted a critical threshold of tumor size at treatment onset, below which ACT^METi^ achieves long-term melanoma control or eradication in line with the experimental data.

**Figure 2.**
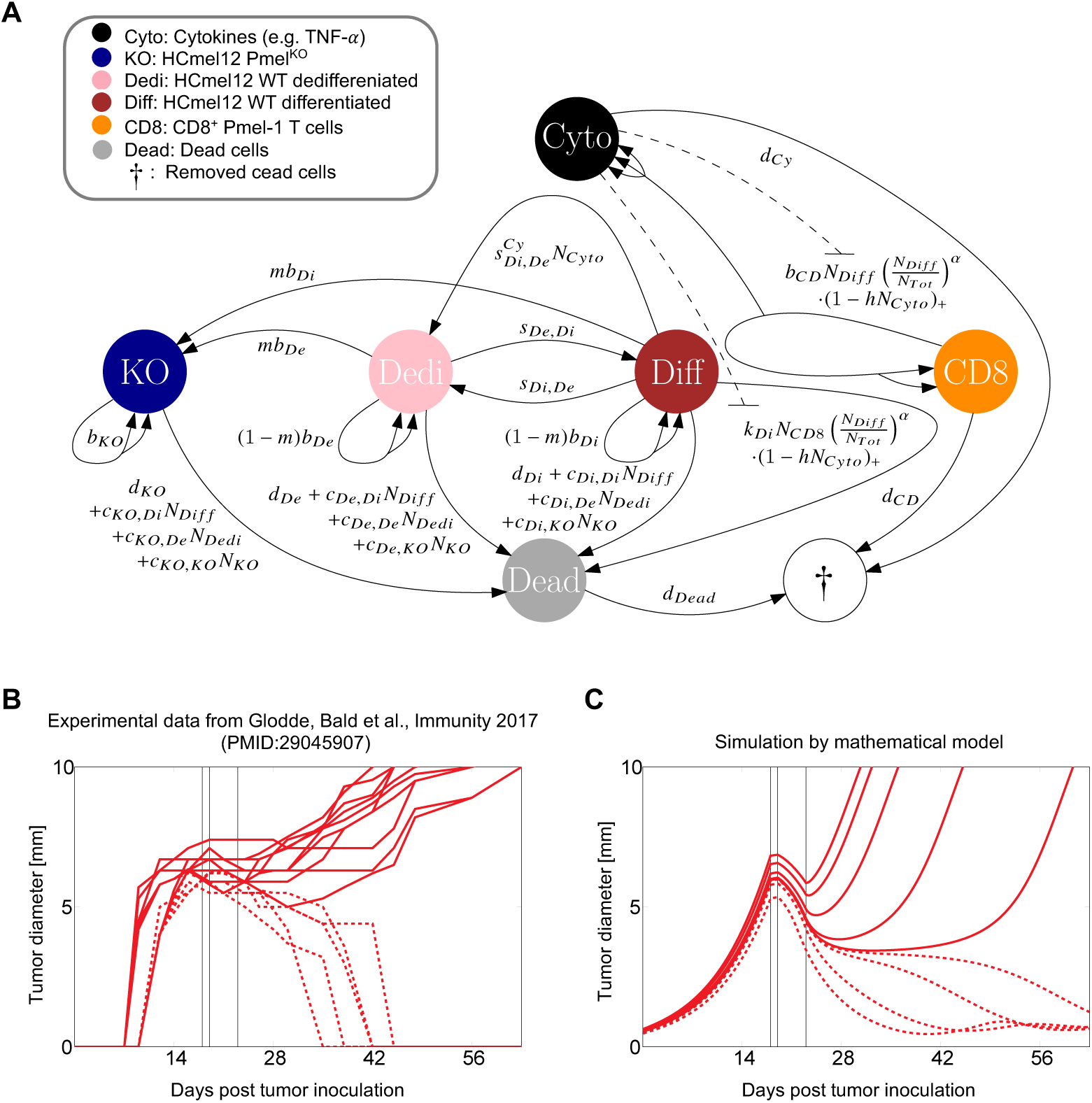
Mathematical model of ACT^METi^ on the basis of an individual-based continuous-time Markov process. **(A)** Interaction diagram displaying cells/molecules (filled circles) and mechanisms that are incorporated in the mathematical model. Arrows represent possible changes to the state of the system, e.g., cell division, cytokine secretion, or cell death. Formulae describe the frequencies at which these population changes occur. **(B)** Tumor growth curves from experimental data published in Glodde et al. **(C)** Tumor growth curves generated by simulations for different initial tumor sizes, shown as tumor diameter [mm]. Vertical lines mark beginning of METi injections, injection of Pmel-1 T cells, and end of METi injections. Dashed lines indicate tumors undergoing eradication.

### Simulating the dynamics of Pmel^KO^ antigen loss variants during ACT^METi^

Next, we simulated the dynamics of pre-existing Pmel^KO^ melanoma cells within a bulk WT melanoma cell population, because we sought to explain the highly variable enrichment of Pmel^KO^ cells found in ACT^METi^-recurrent HCmel12^pcMix^ melanomas from our experiments. In the stochastic model, the context-dependent fitness describes the growth rate of a certain cell type and it is determined by the fixed fitness of its individuals (composed of cell division *b* and death rate *d*) and competition *c* with other individuals. For example, the fixed fitness of Pmel^KO^ cells is r_KO_=b_KO_-d_KO_ while their context-dependent fitness in a population of state N is

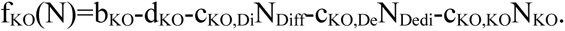

Since the overall growth rate is very similar for untreated pure WT or Pmel^KO^ melanomas (Figure S2C), the reduced fitness of Pmel^KO^ cells in HCmel12^pcMix^ tumors is modeled via a high competitive pressure c_KO,Di_/c_KO,De_ that WT melanoma cells impose on Pmel^KO^ melanoma cells. Figure 3A shows how the growth of the unfit Pmel^KO^ melanoma cell population slows down until a critical number of WT melanoma cells is reached and its overall growth rate (context-dependent fitness) becomes negative. The number of Pmel^KO^ melanoma cells then decreases and would eventually reach zero. In our experiments, however, mice needed to be sacrificed before this would happen.

**Figure 3.**
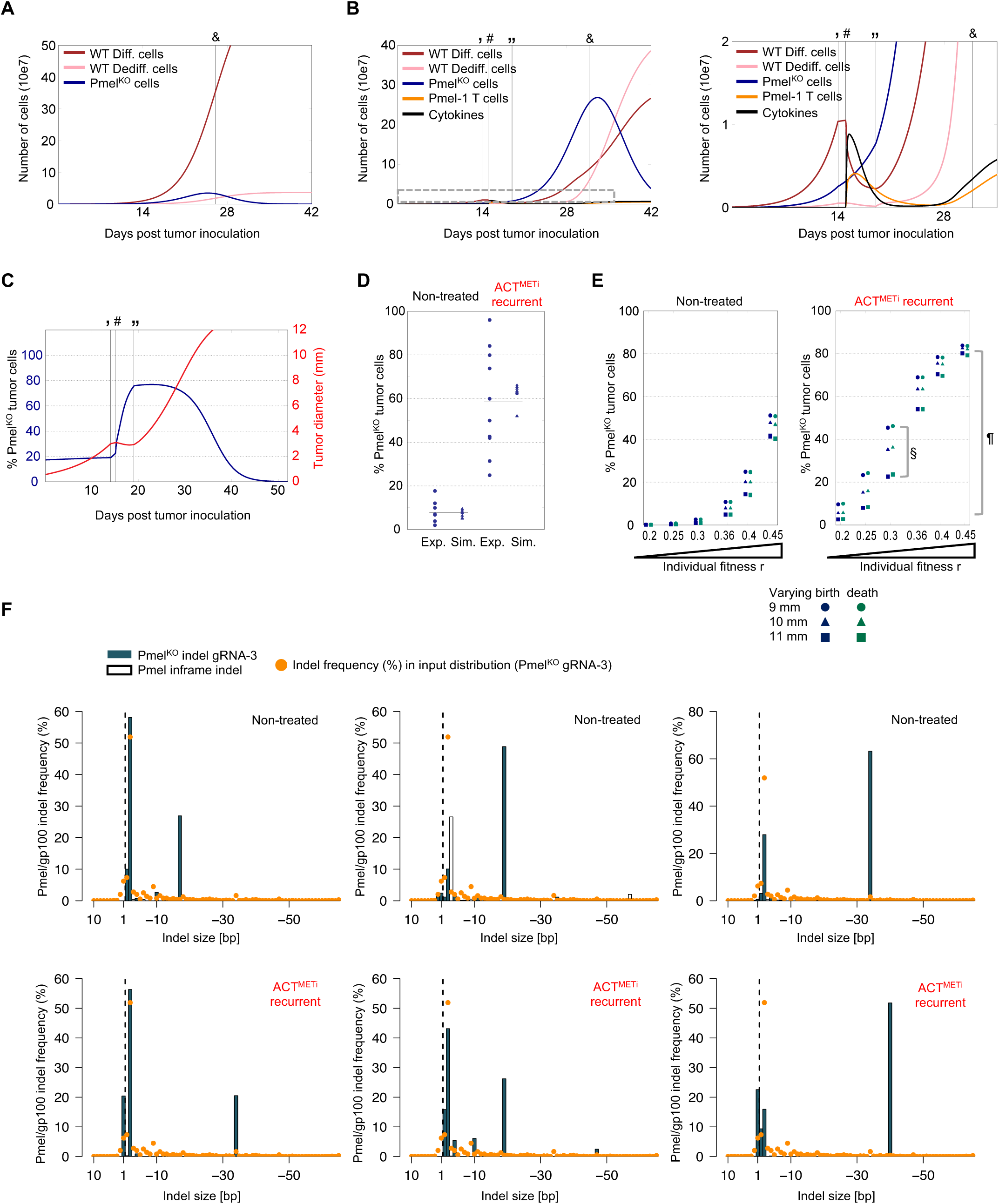
Simulations suggest variable sublcone fitness within the Pmel^KO^ melanoma cell population as cause of highly variable enrichment in ACT^METi^-recurrent tumors. Simulations of the evolution of different cell/molecule types for the untreated case **(A)** and under ACT^METi^ (**B,** left and right zoom-in panel), shown as number of cells in 10e7. **(C)** Simulations of the evolution of tumor size, shown as diameter [mm], and percentage of Pmel^KO^ cells. Initial tumor of medium size and α=4. For A-C vertical lines mark beginning of METi injections (‘), injection of Pmel-1 T-cells (#), and end of METi injections (“) and the time that 10 mm tumor diameter is reached (&). **(D)** Comparison of experimental data (Exp.) from Figure 1G and simulation results (Sim.) for the frequency of Pmel^KO^ alleles in ACT^METi^-recurrent HCmel12^pcMix^ melanomas. Percentage determined at a random time +-0.5 mm of the measured diameter in experiments. **(E)** Predictions for enrichment of Pmel^KO^ cells under varying subclone fitness rKO=bKO-dKO and different tumor sizes at time point of harvesting. Predicted variability of enrichment by different tumor sizes at time point of harvesting (§) verus different sublcone fitness (¶). **(F)** Pmel indel distribution in untreated (upper row) and ACT^METi^-recurrent (lower row) HCmel12^pcMix^ melanomas determined by amplicon NGS. Orange dots indicate respective indel frequency at the time of tumor cell injection (Figure 1D, right panel)

If Pmel-1 T-cell killing of WT^Diff^ melanoma cells was not influenced by the presence of Pmel^KO^ melanoma cells, our simulations predicted that the enrichment of Pmel^KO^ cells in ACT^METi^-recurrent melanomas would be a lot higher than what was measured experimentally, making up nearly the entire melanoma cell population. However, if the portion of Pmel^KO^ melanoma cells is high, one may assume that WT^Diff^ melanoma cells will be ‘shielded’ from Pmel-1 T-cell recognition and killing due to physical obstruction, an immunosuppressive tumor microenvironment and other mechanisms. We model this ‘shielding effect’ by introducing the percentage of WT^Diff^ melanoma cells (out of all tumor cells) as a factor into the killing rate of Pmel-1 T-cells and also their proliferation rate since Pmel-1 T-cell activation depends on Pmel antigen presentation. The influence of this factor is determined by its exponent α, which we termed ‘shielding parameter’. The killing rate thus takes the form

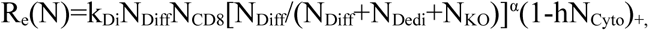

where the last factor models cytokine-induced T-cell exhaustion. The T-cell proliferation rate has the same structure.

A higher value of α causes a larger ‘shielding effect’ through WT^Dediff^ and Pmel^KO^ melanoma cells. Figure 3B shows how, under ACT^METi^, the Pmel^KO^ cell population grows rapidly, as soon as the number of WT cells drops low enough to no longer impose a large competitive pressure (Figure 3B, right zoom-in panel). As a result, the few remaining WT^Diff^ cells are protected by the abundant Pmel^KO^ cells, which allows for the recovery of the WT population. As soon as WT cells surpass the critical level for competition, the number of Pmel^KO^ cells and their proportion of the total melanoma cells decrease again (Figure 3C). Due to these high fluctuations, the measured percentage of Pmel^KO^ cells is highly dependent on (i) the time point of sequencing (harvesting the tumor tissue), and (ii) the influence of the shielding effect. More details on the determination of the parameter α are given in the methods section, supported by Figure S3A and S3B.

### Subclone fitness variability as potential cause of variable immune selection of Pmel^KO^ cells

Figure 3D compares the measured percentage of Pmel^KO^ cells in experiments and simulations, with and without ACT^METi^. The parameters of the mathematical model were chosen such that they match the mean percentage of 7.8% Pmel^KO^ cells in the untreated case and 58.5% under ACT^METi^ (from Figure 1G). Varying the time point of sequencing (harvesting) by picking a random point between 0.5 mm around the documented diameter in the simulations partially explains the fluctuations in the experimental data. Nevertheless, especially in the case of ACT^METi^, we still witness a much higher variation in the experimental data compared to the simulations. Tumor cell heterogeneity is another possibility, which is experimentally modeled in our HCmel12^pcMix^ approach by genetically ablating Pmel antigen expression in thousands instead of raising single cell clones. Each of the generated HCmel12 Pmel^KO^ subclones may display a slightly different fitness due to pre-existing genetic or epigenetic heterogeneity or even CRISPR-Cas9 off-target effects, amongst others. Therefore, we run simulations for varying fitness by changing the birth rate b_KO_ and the natural death rate d_KO_, and thus the overall growth rate. Figure 3E displays the simulated percentage of Pmel^KO^ cells in untreated and recurrent melanomas, determined at tumor sizes between 9 and 11 mm, plotted against their individual fitness r_KO_. The experiments in Figure S2C suggest an average individual fitness of r_KO_=0.36. The results show no major difference between the two approaches of varying b_KO_ and d_KO_. They do, however, account for most of the variation seen in the experimental data as, especially in the case of ACT^METi^ treatment, the percentage of Pmel^KO^ cells at the time point of sequencing (harvesting) largely increases with increasing fitness r_KO_. Thus, in contrast to the tumor size at therapy onset, which surprisingly has very little influence on the Pmel^KO^ cell percentage even when lying above and below the critical threshold for tumor eradication (Figure S3A), varying subclone fitness is a likely explanation for the highly variable enrichment of Pmel^KO^ cells found in ACT^METi^-recurrent melanomas.

### Validating impact of subclone fitness variability on immune selection of Pmel^KO^ cells

To experimentally confirm this prediction from our simulations, we first analyzed the Pmel indel distributions from ACT^METi^-recurrent HCmel12^pcMix^ melanomas. We reasoned that a restriction of the indel distribution, if compared to the input distribution at tumor cell inoculation, would suggest that the population of Pmel^KO^ cells is heterogeneous containing a few Pmel^KO^ subclones with superior fitness that drives their relative enrichment in untreated and ACT^METi^-treated tumors. Indeed, the analysis revealed a few predominant indels deviating from the input indel distribution (Figure 3F). In particular, predominance of indels, which were underrepresented in the input distribution, strongly suggested the selection of a few Pmel^KO^ subclones with superior fitness.

To further substantiate this finding, we revisited our experiments with HCmel12 Pmel^KO^ single cell subclones (Figure S2) now asking whether they exhibited differences in fitness or not. For this purpose, we equally mixed the three Pmel^KO^ subclones together with WT cells to achieve an overall Pmel^KO^ subclone frequency of 25% (8.3% per Pmel^KO^ subclone) (Figure 4A) or 50% (16.7% per Pmel^KO^ subclone) (Figure 4B). Mixed cultures were termed as HCmel12^sccMix^ and injected into the flank of syngeneic mice. Once tumors were established animals were treated with ACT^METi^ or left untreated (Figure 4C-F). Mice were sacrificed when tumors reached the experimental cutoff size of 10 mm in diameter. Frequencies of the three Pmel^KO^ subclones in ACT^METi^-recurrent melanomas were determined by amplicon NGS and we found that clone #1 was predominantly enriched in all cases, whereas enrichment of the other subclones occurred only when the input frequency was increased (Figure 4G). *In vitro,* proliferation of all Pmel^KO^ subclones was comparable to HCmel12 WT cells and if, at all, it was slightly reduced for subclone #1 (Figure 4H). Taken together, selection of HCmel12 Pmel^KO^ single cell clones under ACT^METi^ was strongly biased suggesting substantial differences in the fitness of these subclones.

**Figure 4.**
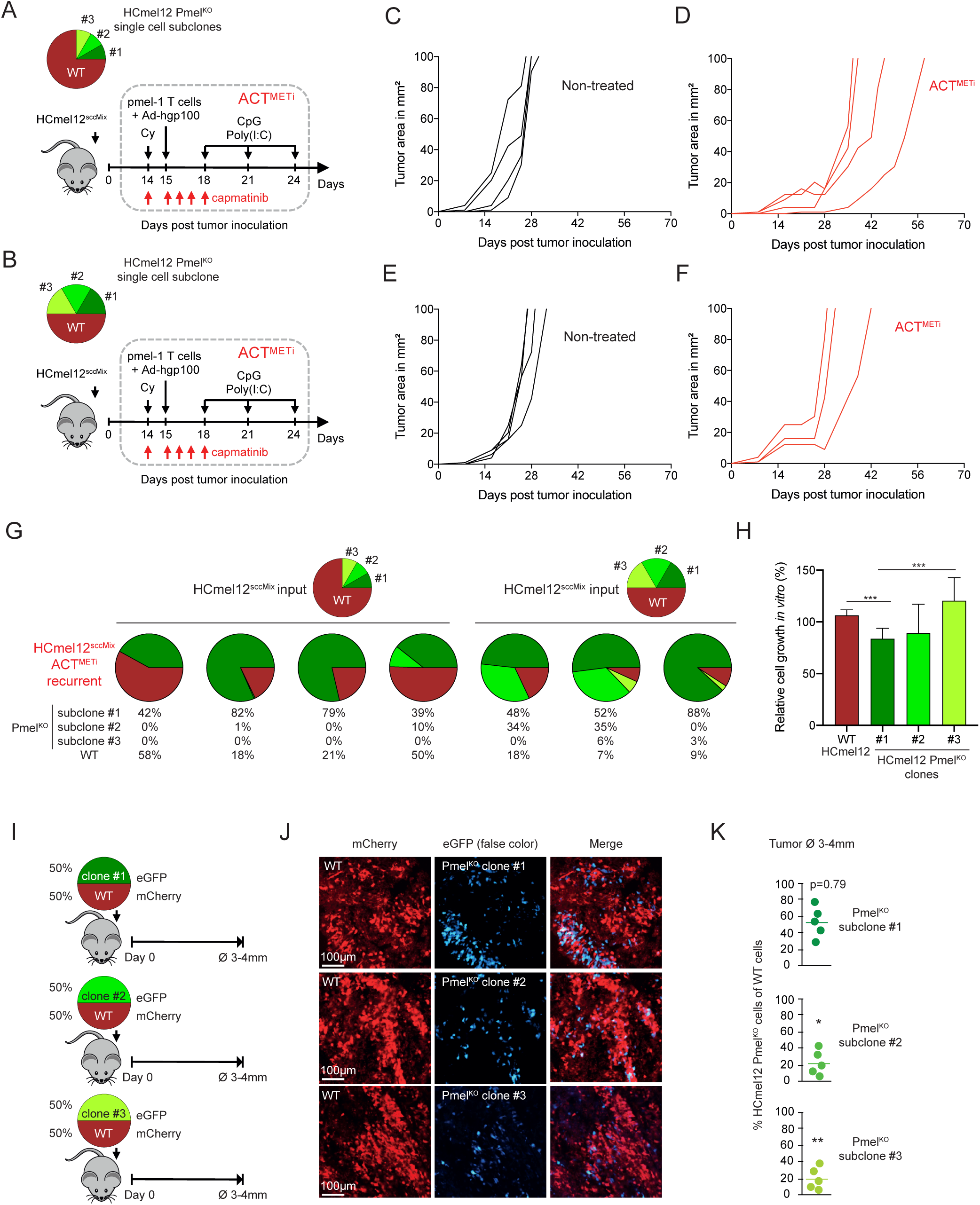
Isolated HCmel12 Pmel^KO^ single cell clones (sublcones) exhibit different fitness in vivo resulting in biased selection under ACT^METi^. **(A-B)** Experimental outline of HCmel12^sscMix^ melanomas treated with ACT^METi^. Mixing ratios of HCmel12 WT and Pmel^KO^ single cell clones are indicated. **(C-F)** Individual tumor growth curves of HCmel12^sscMix^ melanomas described in A-B left untreated or treated with ACT^METi^. **(G)** Percentages of individual Pmel^KO^ single cell clones (subclones) in ACT^METi^-recurrent tumor from D and F determined by amplicon NGS. **(H)** Quantification of in vitro cell growth of individual Pmel^KO^ single cell clones compared to HCmel12 WT cells. Two-sided t test. ***p<0.001. **(I)** Experimental outline of color-coding approach comparing early outgrowth of HCmel12 Pmel^KO^ single cell clones versus WT cells in untreated mice. **(J)** Representative whole-mount fluorescent images corresponding to experimental setup described in I. eGFP signal represented in false color blue. **(K)** Corresponding quantification of J. One sample t-test with hypothetical mean of 50%. *p<0.05, **p<0.01.

To confirm this, we stably transduced HCmel12 WT and the Pmel^KO^ single cell clones with expression constructs encoding for mCherry and tagGFP2, respectively, and injected mixtures at a 1:1 ratio into the flank of syngeneic mice (Figure 4I). Dual-color imaging of whole-mount sections was performed when the tumors had reached a size of 3-4 mm in diameter (Figure 4J). We observed a similar engraftment of HCmel12 WT cells and the Pmel^KO^ single cell clone #1, but the clones #2 and #3 were clearly underrepresented (Figure 4K). Taken together, these results confirmed the prediction from our simulations that Pmel^KO^ subclones substantially differ in fitness providing an explanation for the highly variable enrichment of Pmel^KO^ cells in ACT^METi^-recurrent melanomas.

### Simulating the spontaneous occurrence of Pmel mutations

So far, we only addressed scenarios with pre-existing Pmel^KO^ cells, but if the mutation occurred spontaneously, it would start out with a single cell. In order to determine whether a single Pmel^KO^ cell can fixate although it is unfit compared to the bulk WT cell population, we introduced the possibility of spontaneous mutations from WT cells to Pmel^KO^ cells into our stochastic model. With 1.5 mutations per day on average in a tumor of 3 mm in diameter (typical melanoma size at treatment onset in our experiments), we have chosen a relatively low frequency of mutational events to obtain Figure 5A. For this choice of parameters, in more than half of the simulation runs the Pmel^KO^ cells fixate and cause a relapse within the first 100 days after tumor inoculation. Even when further decreasing the probability of mutation, this still happens, but in less cases and at a later time points. Whenever the Pmel^KO^ cells fixate, i.e. surpass a detectable number of cells, the same effects as in the experimental setup can be witnessed. For a smaller pure WT tumor, below the critical threshold for tumor eradication, the Pmel^KO^ cell population grows and thus protects the WT cells from dying out. The latter can then recover and eventually expand within the Pmel^KO^ tumor. Compared to the situations with a pre-existing portion of Pmel^KO^ cells, this happens much later, because spontaneously occurring Pmel^KO^ cells start to grow from a much lower number. However, the relapse phase itself takes a very similar course (Figure 5A, right zoom-in panel). Figure 5B shows the results of a number of different simulation runs, where the Pmel^KO^ cells fixate at different times to cause a relapse (the occurrence of the mutation that caused the relapse is marked with a cross). This variability is due to the stochasticity of our model where (i) mutations occur randomly at different time points and (ii) Pmel^KO^ mutants may die out before they fixate and hence the first mutation may not always be successful.

**Figure 5.**
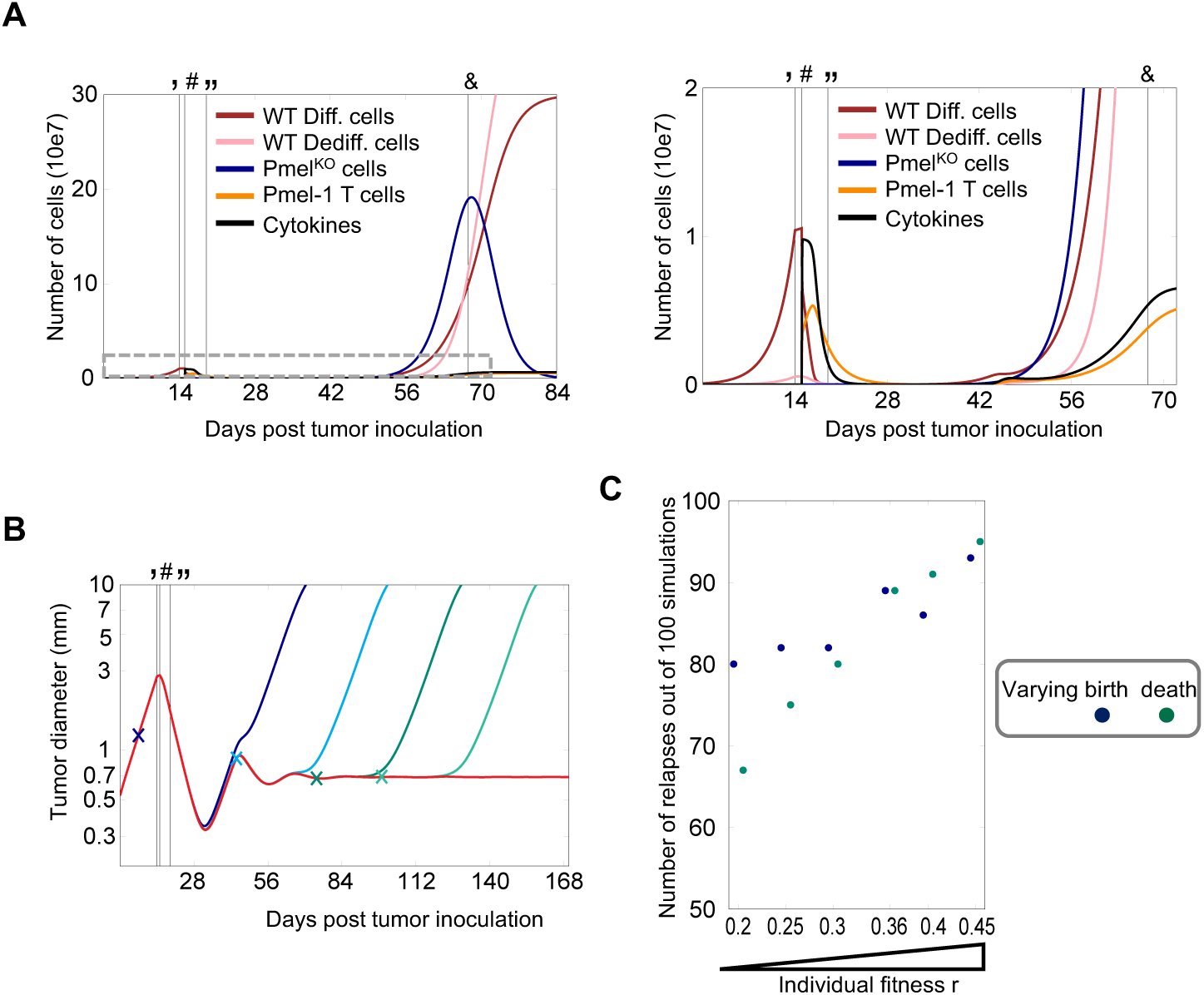
Simulations of spontaneously occurring Pmel^KO^ antigen loss variants suggest cell death inducing as preferred strategy to prevent outgrowth of residual cells. **(A)** Right and left zoom-in panel: Simulation of the evolution of different cell/molecule types under ACT^METi^, shown as number of cells in 10e7. Initial pure WT tumor size below critical threshold for therapy success. Natural mutation to Pmel^KO^ cells at rate of m=10e-7. **(B)** Simulations of tumor growth curves, shown as diameter [mm] on a log-scale. Red curve shows no successful mutation while first successful mutations of blue and green curves are marked with crosses. For A-B vertical lines mark beginning of METi injections (‘), injection of Pmel-1 T-cells (#), and end of METi injections (“) and the time that 10 mm tumor diameter is reached (&). **(C)** Number of tumors going into relapse (reaching diameter of 10 mm) within one year after tumor inoculation, out of 100 runs under varying subclone fitness rKO=bKO-dKO.

### Simulations predict cell death induction as strategy to eliminate residual Pmel^KO^ melanoma cells

As above in Figure 3E, we again varied the individual fitness r_KO_ of the Pmel^KO^ cells. For the percentage of Pmel^KO^ cells at 9 to 11 mm diameter we obtain a similar picture (Figure S4). We see slightly more variability between simulation runs, particularly for high fitness, and on average lower percentages for the intermediate fitness values. However, the overall range in between the highest and lowest values for r_KO_ remains the same and there is no major difference between the variation of b_KO_ and d_KO_. In addition to the percentage of Pmel^KO^ cells, we also studied the likelihood of relapses. Figure 5C shows the number of simulation runs (out of 100) that exhibit a relapse within one year after tumor inoculation. Besides the expectable higher number of relapses for higher fitness r_KO_, we also observe different behavior between the variation of b_KO_ and d_KO_. In the cases where the fitness decrease (compared to r_KO_=0.36) is obtained by an increased death rate d_KO_ or the fitness increase is due to a higher birth rate b_KO_, we detect fewer relapses than in the other cases. This is due to the fact that we have higher rates b_KO_ and d_KO_ (while keeping r_KO_ the same), which causes more birth and death events in the same time interval and thus higher fluctuations in the Pmel^KO^ population. This makes it more likely for the mutant to die out before fixation, i.e. before it reaches a population size at which it cannot go extinct due to random fluctuations, and thus relapses become less likely. Mathematically this is in line with theoretical results calculating the probability of fixation, within a population at state N, to be proportional to f_KO_(N)_+_/b_KO_ and thus to decrease with higher rate b_KO_ (for constant r_KO_ and hence f_KO_(N)). Once a certain population size is reached, the fluctuations have less influence and the Pmel^KO^ cells grow according to their average dynamics, hence their enrichment at relapse is less sensitive to the different approaches. From a clinical perspective, these simulations suggest that active cell death induction is the most effective strategy in order to eliminate residual resistant tumor cells.

## Discussion

We used experimental and mathematical models of melanoma T-cell therapy in order to study the dynamics of antigen loss variants. Tumor cells can downregulate or abolish target antigen expression by epigenetic and genetically hardwired mechanisms (mutations, deletions). In the case of melanoma, melanocyte differentiation antigens like Pmel or MART-1 are well studied targets for antigen-specific T-cell transfer or tumor vaccine therapies both in pre-clinical mouse models and clinical trials (Chodon et al., 2014; Kohlmeyer et al., 2009). However, intratumoral heterogeneous expression patterns and melanoma dedifferentiation, a stress-induced adaptive resistance mechanism, impose challenges for therapeutic efficacy and durability (Landsberg et al., 2012; Mehta et al., 2018). Firstly, our experimental data showed that complete abrogation of Pmel antigen expression by gene editing provided survival benefit to melanoma cells in mice treated with Pmel-specific T-cell therapy (ACT^METi^) despite the fact that melanoma cells can downregulate Pmel expression by dedifferentiation. This result was rather expected, as the importance of high target antigen expression by tumor cells for successful immunotherapy is well established (Leisegang et al., 2016). Of note, the epitope EGSRNQDWL derived from murine Pmel protein is a low affinity epitope (peptide-MHC affinity; IC_50_ 23 μM) and rather poorly recognized by Pmel-1 T cells, inM) and rather poorly recognized by Pmel-1 T cells, in contrast to the corresponding high affinity epitope KVPRNQDWL (peptide-MHC affinity; IC_50_ 186 nM) derived from human PMEL (Engels et al., 2013; Overwijk et al., 1998). Thus, complete loss of antigen expression facilitates tumor immune escape even in the context of a low affinity CD8^+^ T-cell epitope.

To us, it was however unanticipated that the enrichment of engineered Pmel antigen loss variants in recurrent melanomas was subject to such a high variability. In addition, we incidentally found that *Pmel* ablation resulted in a growth disadvantage imposed by wild type melanoma cells when inoculated together in untreated mice. Hence, the fitness of wild type and Pmel^KO^ melanoma cells switched upon ACT^METi^, for which reason we assumed that the immune selection of Pmel^KO^ cells was highly variable because of the dynamics and complexity of interactions between tumor cell variants and T-cells. Mathematical models allowed us to easily manipulate the system and study possible sources for the variable enrichment of Pmel^KO^ cells under ACT^METi^. Given that timing of harvesting tumor tissues for sequencing was standardized in our experimental setup (10 mm tumor diameter), our simulations predicted subclone fitness variability as the most likely cause, which we confirmed by different experimental approaches. We suggest a scenario, where therapy-dependent fitness gain by antigen loss and a variable subclone fitness contribute to a composite fitness that determines subclone dynamics under CD8^+^ T-cell selection pressure. In essence, this underscores the importance of tumor heterogeneity on tumor immune surveillance of melanomas, as also demonstrated by recent studies analyzing melanoma patient samples or using UVB-induced mouse melanomas as a models system (McGranahan et al., 2016; Wolf et al., 2019).

We believe that our findings have important implications for the analysis of patient samples, because the genomic comparison of pre- and post-treatment tumor specimens, untreated and recurrent melanomas in our experimental setting, is a standard approach to identify genetic changes in tumor cells that cause resistance to immunotherapy (Zaretsky et al., 2016). A highly variable enrichment of resistant tumor cell variants limits their detection likelihood, because the chance of being identified as a recurrent event decreases. We therefore postulate that many mechanisms of immunotherapy resistance remain to be discovered, in particular those genetic events that are associated with a reduced tumor cell fitness prior to treatment. Hence, we envision that implementing evolutionary mathematical models into genomic analysis pipelines could help to identify such resistance mechanisms more reliably.

Finally, our simulations of spontaneous antigen loss mutations further emphasized the importance of stochastic modeling, in particular when studying small tumor cell populations. While the deterministic approximation is a good representation of the evolution of large cell populations (Ethier and Kurtz, 1986), random effects are necessary to account for variable relapse times and cause further variation in enrichment of T-cell therapy resistant variants. Moreover, stochastic fluctuations are essential since they can cause extinction of antigen loss variants even though their context-dependent fitness might be positive, i.e., they would grow out in the deterministic model. This ties in with (Champagnat, 2006) calculating the probability of fixation of a mutant to be f_Mut_/ b_Mut_, which, for constant r_Mut_=b_Mut_-d_Mut_, decreases with increasing b_Mut_. In other words, treatments that enforce tumor cell death such as Bcl-2 family antagonists (Ashkenazi et al., 2017) are predicted to efficiently eliminate residual resistant tumor cell variants and thus prevent melanoma recurrence. This scenario reminds of a recent study, where tissue-resident memory CD8 T-cells were shown to achieve long-term immune surveillance of residual melanoma cells in the skin of mice (Park et al., 2019). Experimental models like this seem to be suitable to confirm the prediction regarding cell death induction and may eventually contribute to improved therapeutic strategies that prevent tumor recurrences after successful immunotherapy.

## Author contributions

Conceptualization, T.T., A.B., M.H.; Methodology, N.G., D.vdBK., T.B.; Algorithms, A.K., K.E., F.K., M.R., JL.SB, A.B.; Software, A.K., K.E., M.R., JL.SB; Validation, N.G., D.vdBK., S.V., P.A., T.B.; Formal Analysis, N.G., A.K., T.B., J.L., M.H.; Investigation, N.G., D.vdBK., S.V., T.B., P.A.; Resources, N.G., T.B., J.L., T.T., M.H.; Data Curation, N.G., A.K., T.B., M.H.; Writing – Original Draft, N.G., A.K., A.B., M.H.; Writing – Review & Editing, T.T., A.B., M.H.; Supervision, M.R., J.L., A.B., T.T., M.H.; Project Administration, T.T., A.B., M.H.; Funding Acquisition, T.T., A.B., M.H. The authors declare no conflict of interest.

## Acknowledgements

We thank P. Wurst and An. Dolf from the UKB FACS core facility for help with flow cytometry and cell sorting. T.T. was funded in part by grants from the DFG (TU 90/8-1 and A27 in the SFB854). A.B. and M.R. were funded by the Deutsche Forschungsgemeinschaft (DFG, German Research Foundation) under Germany’s Excellence Strategy - GZ 2047/1, Project-ID 390685813. A.B. and M.H. were supported by the Deutsche Forschungsgemeinschaft (DFG, German Research Foundation) under Germany’s Excellence Strategy – EXC2151 – Project-ID 390873048. T.T. and M.H. were funded by Else-Kröner-Fresenius-Stiftung (EKFS 2013_A297).

## METHODS

## CONTACT FOR REAGENT AND RESOURCE SHARING

Further information and requests for resources and reagents should be directed to and will be fulfilled by the Lead Contact, Michael Hölzel. Request regarding source code (C++) of the mathematical model will be fulfilled by.

## EXPERIMENTAL MODEL AND SUBJECT DETAILS

### Mice

C57BL/6J mice (H2-Db) were purchased from Charles River. T-cell receptor (TCR)-transgenic Pmel-1 mice expressing an ab TCR specific for amino acids 25-33 of human (high affinity) and mouse (low affinity) Pmel (gp100) presented by H2-Db were bred as described previously (Engels et al., 2013; Kohlmeyer et al., 2009; Landsberg et al., 2012; Overwijk et al., 2003). All animals were maintained under specific pathogen-free conditions in individually ventilated cages and experiments were performed with 6-8 weeks old mice. All experiments were approved by the responsible authorities (Landesverwaltungsamt, SA, and LANUV, NRW; Germany) and were performed according to the institutional and national guidelines for the care and use of laboratory animals.

### Cell line

The melanoma cell line HCmel12 was established from the Hgf-Cdk4^R24C^ melanoma model as described previously (Bald et al., 2014). Monoclonal and polyclonal HCmel12 Pmel^KO^ variants were generated using the CRISPR-Cas9 genome engineering technology (see below in the “method details” section). All HCmel12 melanoma cell variants were routinely cultured in “complete RPMI medium”, i.e. RPMI 1640 medium (Life Technologies) supplemented with 10% FCS (Biochrome), 2 mM L-glutamine, 10 mM non-essential amino acids, 1 mM Hepes (all from Life Technologies), 20 µM 2-mercaptoethanol (Sigma), 100 IU/ml penicillin and 100 µg/ml streptomycin (Invitrogen). All cell lines used in this study were routinely tested for mycoplasma contamination by PCR.

## METHOD DETAILS

### Generation of gp100 (Pmel/silv) sgRNA CRISPR-Cas9 plasmids

px330-U6-Chimeric_BB-CBh-hSpCas9 (px330) (Addgene, plasmid #42230) was digested with BbsI and gel purified. Small DNA oligonucleotides (Microsynth) representing sgRNAs against exon 1 of murine *Pmel* (also known as gp100) were annealed and cloned into digested pX330. The following target sequences were used (underlined PAM sequences NGG at the end were not included in the oligo) based on design rules described at http://www.genome-engineering.org/):

sgRNA#1, GCTTGTGCTGAGTGCTCTGC<u>TGG</u>;

sgRNA#2, GCTGAGTGCTCTGCTGGCTG<u>TGG</u>;

sgRNA#3, GAGTGCTCTGCTGGCTGTGG<u>GGG</u>.

3 μM) and rather poorly recognized by Pmel-1 T cells, ing from each oligo was mixed to annealing buffer (100mM NaCl and 50mM Hepes pH7.4) in 50 μM) and rather poorly recognized by Pmel-1 T cells, inl total volume. This mixture was incubated for 4 minutes at 90°C, then 10 minutes at 70°C. Next, the annealed oligos were slowly cooled down to 10°C. 2μM) and rather poorly recognized by Pmel-1 T cells, inl of the annealed oligos were ligated into 100 ng of linearized vector (pX330). The ligated product was then transformed into DH10ß chemo-competent bacteria and correct clones were identified by sequencing.

### Generation of HCmel12 Pmel^KO^ cultures

5*10^5^ HCmel12 melanoma cells were seeded in 12-well plates 2 hours prior to reverse transfection. The cells were transfected with 2μM) and rather poorly recognized by Pmel-1 T cells, ing plasmid (mix of 1.6μM) and rather poorly recognized by Pmel-1 T cells, ing px330-sgRNA and 0.4μM) and rather poorly recognized by Pmel-1 T cells, ing pRp-GFP) using Fugene HD transfection reagent (Promega) according to the manufacturer’s protocol. DNA:Fugene HD ratio of 1:3 was determined to be the optimal condition for HCmel12 cells. GFP positive cells were sorted (BD FACSAria II Cell Sorter; BD Biosciences) after 48 hours and expanded. For generation of gene-edited Hcmel Pmel^KO^ single cell clones, cells were seeded at 0.7 cells per 96-well. Single cell clones were further expanded until a 70 percent confluence was achieved. Single cell clones with successfully targeted *Pmel* gene were identified by amplicon next generation sequencing (Illumina MiSeq platform). For polyclonal Pmel gene editing approaches, the frequency of Pmel frame-shift and in-frame indels of FACS sorted cultures was determined by amplicon next generation sequencing and a modified version of the OutKnocker webtool (http://www.outknocker.org) (Schmid-Burgk et al., 2014).

### Amplicon next generation sequencing

Genomic DNA (gDNA) from cultured cells and homogenized melanoma tissues was extracted using the Nucleo Spin Tissue kit (Macherey&Nagel) according to the manufacturer’s recommendations. For the generation of targeted PCR amplicons for next generation sequencing (NGS), a two-step PCR protocol was performed. For the first PCR, a *Pmel* gene-specific primer pair (Pmel_Ex1_fwd & Pmel_Ex1_bwd; encompassing the sgRNA target site was used with additional adapter sequences for the second PCR. The *Pmel* gene-specific sequences are underlined:

Pmel_Ex1_fwd, ACACTCTTTCCCTACACGACGCTCTTCCGATCT<u>AAGGCCTATGCAAATGACCATC;</u>

Pmel_Ex1_bwd, TGACTGGAGTTCAGACGTGTGCTCTTCCGATCT<u>AGGACAAGCCTAAAGTATTTTGACTTG.</u>

Adapter-specific universal primers containing barcode sequences and the Illumina adapter sequences P5 and P7 were used for the second PCR (Combinations of D501.. and D701.. primers). In the first PCR, the genomic region of interest was amplified with 18 cycles, using approximately 20-50 ng of gDNA as input and Phusion HD polymerase (New England Biolabs) in a 12.5 μM) and rather poorly recognized by Pmel-1 T cells, inl mixture according to manufacturer’s protocol. Next, 2 μM) and rather poorly recognized by Pmel-1 T cells, inl were transferred to the second PCR. This product was amplified with another 18 cycles in a 25 μM) and rather poorly recognized by Pmel-1 T cells, inl reaction mix with Phusion HD polymerase. Next-generation sequencing was performed with MiSeq Gene & Small Genome Sequencer (Illumina) according to manufacturer’s standard protocols with a single-end read and 300 cycles (MiSeq Reagent Kit v2 300 cycle).

### Tumor transplantation experiments

Cohorts of syngeneic C57BL/6 mice were injected intracutaneously (i.c.) with a total of 2*10^5^ HCmel12 melanoma cells (WT, Pmel^KO^ or mixtures thereof as described in the main text) into the flank. Polyclonal HCmel12 Pmel^indel^ cultures (#1, #2 and #3) were mixed with parental HCmel12 (HCmel12 WT) cells in order to obtain a Pmel indel frequency of 20%. For monoclonal approaches, three independent HCmel12 Pmel^KO^ clones (#1, #2 and #3) were mixed in the same ratio and injected either alone or mixed with HCmel12 WT cells to achieve an overall Pmel^KO^ single cell clone frequency of 25 or 50%, respectively. Tumor size was measured twice weekly and recorded as diameter in millimeter. Mice with tumors exceeding 10 mm in diameter or signs of illness were sacrificed for animal welfare reasons.

### Multimodal adoptive T-cell immunotherapy (ACT) plus c-MET inhibition

Adoptive T cell transfer immunotherapy (ACT) was performed as previously described (Glodde et al., 2017; Kohlmeyer et al., 2009; Landsberg et al., 2012). In brief, when transplanted tumors reached a size of 3-5 mm in diameter mice were preconditioned for ACT by intraperitoneal (i.p.) injection of 2 mg (100 mg/kg) cyclophosphamide 24 hours before adoptive transfer of Pmel-1 T cells. One day later, 2*10^6^ naïve Pmel-specific TCR transgenic CD90.1^+^CD8^+^Vß13^+^ T-cells, isolated from spleens of Pmel-1 transgenic mice were intravenously (i.v.) injected and activated *in vivo* by a single intraperitoneal dose (5*10^8^ PFU) of the recombinant adenoviral vector Ad-gp100 (Kohlmeyer et al., 2009). On days 3, 6 and 9 after adoptive pmel-1 T cell transfer, mice received 50µg of CpG 1826 DNA (MWG Biotech) and 50 µg of polyinosinic:polycytidylic acid (Invivogen) in 100µl of PBS peritumorally. The complete ACT protocol was combined with intraperitoneal injections of a c-MET inhibitor (5 mg/kg KG METi, capmatinib; Novartis and Selleck Chemicals) in 100 µl PBS every 12 hours for 5 consecutive days as previously described (Glodde et al., 2017).

### Histology and immunohistology

Tumor tissue samples were immersed in a zinc-based fixative (BD Pharmingen). Tissues were embedded in paraffin and sections stained with rabbit anti-mouse Pmel pAb (NBP1-69571; Novus Biologicals) followed by enzyme-conjugated secondary antibodies and the LSAB-2 color development system (DAKO). Heavily pigmented mouse melanomas were bleached before staining (20 min at 37°C in 30% H_2_O_2_ and 0.5% KOH, 20 sec in 1% acetic acid and 5 min in TRIS buffer). Stained sections were examined with a Zeiss Axio Observer Z1 microscope. Images were acquired with a Zeiss AxioCam ICc5 digital camera and processed with Adobe Photoshop. The percentage of Pmel (gp100)-positive melanoma cells was evaluated in five high-power fields per tumor by three independent investigators.

### Flow cytometry

To analyze the effect of c-MET inhibition on inflammation-induced dedifferentiation *in vitro*, HCmel12 and B16F1 melanoma cells were stimulated with recombinant murine TNF-α (1000 U/ml; Peprotech) or 100 nM capmatinib (Selleck Chemicals) alone or in combination for 72 hours. Vehicle treated cells served as controls. Cells were stained with a biotinylated polyclonal antibody specific for mouse NGFR (R&D) followed by an appropriate Streptavidin conjugated secondary antibody according to standard protocols. Data were acquired with a FACSCanto flow cytometer (BD Biosciences) and analyzed with the FlowJo software (TreeStar, V7.6.5 for Windows).

### Determining HCmel12 tumor cell content

To determine the HCmel12 tumor cell content in whole-tumor DNA preparations, the genomic region containing the HCmel12 private p53^R172H^ mutation was amplified by a two-step PCR. First we used gene-specific primers followed by barcoded primers (Illumina barcodes: D501-508 & D701-D712) containing the Illumina adapters P5 and P7. The following primers were used for the p53 gene (*Trp53*) specific first PCR (gene specific sequences underlined):

Trp53_NG1_fwd, ACACTCTTTCCCTACACGACGctcttccgatct<u>CCCTCAATAAGCTATTCTGCCAG</u>;

Trp53_NG1_bwd, TGACTGGAGTTCAGACGTGTGCTCTTCCGATCT<u>AGGCCTAAGAGCAAGAATAAGTC.</u>

Pooled PCR products were sequenced on the Illumina MiSeq platform. We used the R-based Bioconductor programming environment to import raw FASTQ files and performed the read alignment to the genomic region of mouse p53 with the align function of the Rsubread package. The deepSNV package was used to determine p53^wt^ versus p53^R172H^ ratios at the position encoding for the p53^R172H^ mutation. Pmel knockout allele frequencies were divided by the tumor cell content (mutant p53 allele ratio) for normalization.

### Indel detection

For basic indel detection we used the web-based program outknocker (http://www.outknocker.org/) or a slightly modified version thereof (Schmid-Burgk et al., 2014). FASTQ files were imported and the sequence of the Pmel PCR amplicon was used as reference sequence for alignment. To determine polyclonal indel distribution and quantify frame-shift and in-frame indels, individual reads were transferred into an indel-size/indel-position matrix. This analysis was done for polyclonal genome-edited HCmel12 cultures and untreated versus ACT^METi^-treated HCmel12 tumors. All analyses were done using the R-based statistical computing platform.

### Generation of HCmel12^METi-R^ cells

Spontaneous METi-resistant HCmel12 cells (HCmel12^METi-R^) were raised by continuous exposure of low-density seeded parental HCmel12 cells with the c-MET inhibitor capmatinib (100 nM). Three independent colonies arising approximately after three weeks of continuous exposure were isolated, propagated and further characterized by colony formation assays, Western blot analysis and gene expression microarray (data not shown). For a single round of *in vivo* passaging, 2*10^5^ HCmel12^METi-R^ cells were injected into the flank of syngeneic C57BL/6J mice and *ex vivo* HCmel12^METi-R^ cultures were established after sacrificing the mice.

### Cell growth assays

3000 HCmel12 melanoma cell variants were plated in 12-well plates and treated with the MET inhibitor capmatinib (also known as INC280) or the MEK inhibitor trametinib (also known as GSK1120212) (all inhibitors for *in vitro* assays from Selleck Chemicals) at indicated concentrations or vehicle control for 6 days. Dishes were stained with a standard crystal violet staining procedure. In brief, cells were washed with PBS, fixed in 4% formaldehyde solution and stained with 0.05% crystal violet in water for 30min. Stained dishes were washed three times with water to remove background staining. Colonies were scanned and quantified using the Odyssey SA Infrared Imaging System (LICOR Biosciences) as surrogate measure for cell number.

### Immunoblots

Protein lysates were prepared from cultured cells after 2 or 72 hours of incubation with capmatinib (INC280), trametinib (GSK1120212) or vehicle control and lysed directly in Laemmli buffer (SDS loading buffer). The lysates were incubated for 5 minutes at 95°C prior to loading. Samples were loaded and separated by SDS–PAGE gel electrophoresis and transferred to nitrocellulose membrane with 0.2 μM) and rather poorly recognized by Pmel-1 T cells, inm pore size (GE Healthcare) according to standard protocols. Blots were immunostained with p44/42 MAPK (ERK1/2) rabbit monoclonal antibody (#9102; Cell Signaling), phospho-ERK (E-4) mouse monoclonal antibody (sc-7383; Santa Cruz), Akt (pan) (40D4) mouse monoclonal antibody (#2920; Cell Signaling), phospho-Akt (Ser473) (D9E) rabbit monoclonal antibody (#4060; Cell Signaling), ß-Actin (C4) mouse monoclonal antibody (sc-47778; Santa Cruz), gp100/PMEL/SILV polyclonal goat antibody (NB100-41098; Novus Biologicals), c-Met (B-2) mouse monoclonal antibody (sc-8057; Santa Cruz) and phospho-Met (Tyr1234/1235) (D26) rabbit monoclonal antibody (#3077; Cell Signaling). Bound antibodies were visualized with IRDye 680RD or IRDye 800CW secondary antibodies (LICOR Biosciences) for detection in the 700 nm and 800 nm channel, respectively. Blots were scanned with the Odyssey SA Infrared Imaging System (LICOR Biosciences).

### Analysis of spatial growth patterns in transplanted melanoma by genotype-dependent color-coding

For whole mount immunofluorescence analysis melanoma cells were transduced with retroviral vectors generated using the proviral constructs pRp-tagGFP2 or pRp-mCherry and two retroviral packaging plasmids pMD.2G (expressing VSVg) and pCMVGag-Pol as described (Riesenberg et al., 2015). mCherry-expressing HCmel12 WT and tag-GFP2 expressing Pmel^KO^ single cell clones were individually mixed at a ratio 1:1 and injected intracutaneously into the flank of C57BL/6 mice. Mouse skin melanomas with a diameter of 3-4 mm were harvested, fixed in PLP-Buffer (0.05 M phosphate buffer containing 0.1 M L-lysine [pH 7.4], 2 mg/ml NaIO4, and 10 mg/ml paraformaldehyde) over night at 4°C and mounted with Fluoromount-G (Southern Biotech). Images were acquired with an upright LSM780 confocal laser-scanning microscope (Carl Zeiss Microimaging) equipped with a Plan Apochromat 20x/0.8. Image analysis was performed using the Imaris software (Bitplane) as described previously (Bald et al., 2014; Glodde et al., 2017).

## QUANTIFICATION AND STATISTICAL ANALYSIS

### Statistical analyses

Statistical significance of experimental results was evaluated with GraphPad Prism 8 software or R computing platform using the parametric unpaired two-tailed student’s t-test or the non-parametric Wilcoxon-Mann-Whitney test depending of the type of source data (e.g. normal distribution). The use of statistical tests is specified in the figure legends. P-values less than 0.05 were considered significant (*p<0.05, **p<0.01, ***p<0.001).

## MATHEMATICAL MODELING

## UNDERLYING MODEL

### Stochastic model

To study the evolution of melanomas under the improved ACT^METi^ therapy, we use an individual-based continuous-time Markov process that is an extension of the model in (Baar et al., 2016). This model itself is based on an individual-based model of adaptive dynamics, introduced in (Bolker and Pacala, 1997, 1999; Dieckmann and Law, 1996; Law and Dieckmann, 1999; Fournier and Méléard, 2004) and further developed over the last years (Champagnat, 2006; Champagnat and Méléard, 2011; Champagnat et al., 2008), in particular for the scenario of phenotypic switching (Baar and Bovier, 2018). The tumor is composed of a finite number of different cells and cytokines. The state of the system, i.e. the number of different cells/molecules at time t, is described by the vector N(t)=(N_Diff_(t),N_Dedi_(t),N_KO_(t),N_Dead_(t),N_CD8_(t),N_Cyto_(t)), where we distinguish differentiated, dedifferentiated, Pmel^KO^, and dead (not yet disintegrated) melanoma cells, as well as antigen-specific cytotoxic CD8+ Pmel-1 T-cells (for simplicity abbreviated as T-cells in the following section), and cytokines (e.g. TNF-α and IFN-γ). The dynamics of the system are determined by a number of events that change the state of the population and occur at certain rates or frequencies. Those rates determine the exponential waiting time until the next event occurs and depend on fixed parameters as well as the current state of the population. The evolution of the Markov process is described by its infinitesimal generator that is of the form

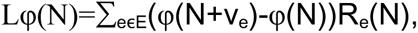

where E is the set of possible events, v_e_ is the change in the population associated to an event, R_e_(N) is the rate at which the event occurs, and φ is a measurable bounded function. It can be constructed rigorously similar to the process in (Fournier and Méléard, 2004).

In our model we consider the following events and rates (visualized in Figure 2A):

Differentiated WT melanoma cells

– reproduce clonal at rate (1-m)b_Di_N_Diff_
– switch to dedifferentiated state (naturally) at rate s_Di,De_N_Diff_
– die (naturally and due to competitive pressure) at rate (d_Di_+c_Di,Di_N_Diff_+c_Di,De_N_Dedi_+c_Di,KO_N_KO_)N_Diff_

Dedifferentiated WT melanoma cells

– reproduce clonal at rate (1-m)b_De_N_Dedi_
– switch to differentiated state (naturally) at rate s_De,Di_N_Dedi_
– die (naturally and due to competitive pressure) at rate (d_De_+c_De,Di_N_Diff_+c_De,De_N_Dedi_+c_De,KO_N_KO_)N_Dedi_

Pmel^KO^ melanoma cells

– reproduce clonal at rate b_KO_N_KO_
– arise as mutants from (de)differentiated melanoma cells at rate m(b_Di_N_Diff_+b_De_N_Dedi_)
– die (naturally and due to competitive pressure) at rate (d_KO_+c_KO,Di_N_Diff_+c_KO,De_N_Dedi_+c_KO,KO_N_KO_)N_KO_

Dead melanoma cells

– get disintegrated at rate d_Dead_N_Dead_

CD8 Pmel-1 T-cells (abbreviated as T-cells)

– reproduce/get activated/get recruited and simultaneously secrete l cytokines at rate b_CD_N_Diff_N_CD8_[N_Diff_/(N_Diff_+N_Dedi_+N_KO_)]^α^(1-hN_Cyto_)_+_
– kill differentiated melanoma cells at rate k_Di_N_Diff_N_CD8_[N_Diff_/(N_Diff_+N_Dedi_+N_KO_)]^α^(1-hN_Cyto_)_+_
– die/get inactive at rate d_CD_N_CD8_

Cytokines

– induce an additional dedifferentiation of melanoma cells at rate s^Cy^_Di,De_N_Cyto_N_Diff_
– get disintegrated at rate d_Cy_N_Cyto_

Most rates correspond to a standard birth-and-death-model with competition and switching/mutation. The unusual rates are the ones of T-cell reproduction and melanoma cell killing. They do not only depend on the number of differentiated melanoma cells and T-cells but include two additional factors. [N_Diff_/(N_Diff_+N_Dedi_+N_KO_)]^α^ represents the effect of differentiated cells being shielded from the T cells by other melanoma cells, not susceptible to the T-cells. It partially takes into account the spatial structure of the tumor. The proportion of differentiated cells [N_Diff_/(N_Diff_+N_Dedi_+N_KO_)] is weighed by a parameter α that determines the influence of this shielding effect. The factor (1-hN_Cyto_)_+_ corresponds to cytokine-induced T-cell exhaustion. As the number of inflammatory cytokines increases, the melanoma cells up-regulate PD-1 ligands (PD-L1) that inhibit the immune reaction.

The injections of METi are not modeled by an additional particle, but as a change in the parameters during the five days of injections. This is reasonable since regular injections ensure a relatively stable level of METi, which is then degraded quickly after the injections stop. Melanoma cell reproduction and T cell exhaustion are down-regulated, while reproduction and killing activity of T-cells are up-regulated.

Dead melanoma cells are included in the model since they contribute to the experimentally measured diameter of the tumor.

### Deterministic approximation and hybrid algorithm

In simulations, we use a Gillespie-type algorithm that generates a realization of the stochastic process by simulating single events (Gillespie, 1976). Since this method is computationally intensive in large populations with frequent events, we combine stochastic simulations of rare events and deterministic simulations (employing Runge-Kutta methods (Butcher, 1963; Kutta, 1901; Runge, 1895)) of frequent events to reduce the running time while keeping random effects such as subpopulations dying out. Similar approaches have been discussed in (Marchetti et al., 2016; Salis and Kaznessis, 2005). This is reasonable due to an approximation result by Ethier and Kurtz (Ethier and Kurtz, 1986): For a large system, where the number of particles (cells, molecules etc) is of order K, the relative number of particles N(t)/K behaves approximately like the solution to a deterministic system of differential equations. If, for each event e, the population changes from N to N+v_e_ at rate K*R’_e_(N/K), then N(t)/K≈n(t) with

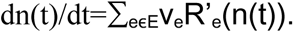

## PARAMETER CHOICES

### Derivation from experiments

We have two sets of parameters corresponding to two sets of experiments. First, the experiments with pure wild type tumors from Glodde et al. (Glodde et al., 2017), which are displayed in Figure 2B, and second, the experiments including Pmel^KO^ cells that are new to this paper. Between the sets of experiments, the melanoma cells showed slightly different behavior (e.g. different speed of growth), however this can be achieved through variation of the parameters, leaving the systemic level (different events, structure of the rates) unchanged.

We assume that the tumor takes the form of a 3-dimensional ball to relate the number of cells in our model to the tumor diameter measured in the experiments. A 5 mm tumor contains roughly 7*10e7 cells and since the measured tumor size varies from 1 mm ≈ 5.6*10e5 cells to 10 mm ≈ 5.6*10e8 cells, we choose K=10e7 as the typical size for the deterministic approximation.

The number of initially injected melanoma cells is 2*10e5. Since likely not all of these cells contribute to form the growing tumor, the initial condition N_Diff_(0)+N_Dedi_(0)+N_KO_(0) is varied between 10e4 and 2*10e5.

The growth parameters of the different melanoma cells can be approximated from experiments where tumor cells are injected into mice and then left to grow without treatment. We set the death rates to 0.1 and assume the same rates for differentiated and dedifferentiated wild-type tumor cells, since we cannot distinguish them in the experiment. Through a logistic fitting, we determine the birth and self-competition rates. The cross-competition between Pmel^KO^ and wild-type cells is only relevant to the new experiments and is chosen to fit the percentages in Figure 1G (results in Figure 3D).

The rates for natural and cytokine-induced switching between differentiated and dedifferentiated wild-type melanoma cells are chosen to fit experiments from Figure S1F, where wild-type melanoma cells are treated in vitro with TNF-α and/or METi. Without treatment, a ratio of 0.95:0.05 of differentiated to dedifferentiated melanoma cells is observed, which determines the ratio between the natural switch rates. This ratio shifts to 0.65:0.35 under the influence of TNF-α, which indicates an additional, cytokine-induced, dedifferentiation. Under addition of METi alone, the ratio is the same as in the untreated case, while a combination with TNF-α results in a ratio of 0.85:0.15. This suggests that METi has no influence on the natural switch rates, while partially cancelling the cytokine-induced switch.

The therapy parameters (reproduction, death, killing efficiency, and exhaustion of T-cells, secretion and degradation of cytokines) are chosen to fit the experiments in Figure 2B and experiments with pure wild-type tumors corresponding to the protocol in Figure 1A. During the course of therapy, 2*10e5 specific T-cells are injected into the mice, which do not all infiltrate the tumor tissue. Simulations show that a variation of the number of T-cells has little to no effect on the evolution of the tumor, therefore we fix it to 10e5 for all simulation runs. The variation between different mice is obtained by running simulations with different initial conditions (initial number of melanoma cells). The effect of METi on the proliferation, killing efficiency, and exhaustion of T-cells is determined comparing experiments with therapy protocols including and excluding METi injections.

The parameter for the degradation of dead melanoma cells is chosen to fit the descent of tumor size during the 5 days of METi injections. During this phase, the most dead melanoma cells are produced due to effective killing by T cells, which contributes to the measured tumor diameter.

The parameter α, which scales the effect of differentiated cells being shielded from T-cells by other melanoma cells, is chosen to fit the percentages in Figure 1G (reseults in Figure M3D). The parameter has little influence in the simulations with pure wildtype tumors, which is why we determine it after fitting the other therapy parameters. An increase in α corresponds to a larger shielding effect. Since the experimental data in Figure 1G has a broad spectrum, we investigate several sources of variation to determine α. In Figure S3A, the composition of the tumor is analysed at a diameter of 10 mm, while the initial conditions are varied (keeping the initial percentage of Pmel^KO^ cells at 17,1%). Even though we compare initial sizes that lie below and above the critical threshold for therapy success, the percentages are very similar. Therefore, we fix the initial number of cells to a medium amount of 10e5 cells in Figure S3B and vary the time at which the percentage is taken between 9 and 11 mm diameter. Figure 3C shows that the choice of this time has a big influence on the percentage of Pmel^KO^ cells. Temporarily, the those cells make up a large portion of the tumor with up to 80%, before the wild-type cells recover and the context-dependent fitness of Pmel^KO^ cells become negative again due to the increased competitive pressure. To obtain an average of around 60%, we set α to 4.

|  | Glodde et al.,<br>ACT | Glodde et al.,<br>ACT<sup>METi</sup> | This paper,<br>ACT | This paper,<br>ACT<sup>METi</sup> |
| --- | --- | --- | --- | --- |
| K | 10e7 | 10e7 | 10e7 | 10e7 |
| $b_{Di}=b_{De}$ | 0.5 | 0.113 | 0.45 | 0.113 |
| $b_{KO}$ | - | - | 0.46 | 0.33 |
| $d_{Di}=d_{De}=d_{KO}$ | 0.1 | 0.1 | 0.1 | 0.1 |
| $c_{Di,Di}=c_{Di,De}=c_{De,Di}=c_{De,De}$ | 0.0053 | 0.0002 | 0.0047 | 0.0002 |
| $c_{KO,KO}$ | - | - | 0.0048 | 0.003 |
| $c_{Di,KO}=c_{De,KO}$ | - | - | 0.00002 | 0.00002 |
| $c_{KO,Di}=c_{KO,De}$ | - | - | 0.0117 | 0.0117 |
| $d_{Dead}$ | 0.5 | 0.5 | 0.5 | 0.5 |
| $s_{Di,De}$ | 0.05 | 0.05 | 0.05 | 0.05 |
| $s_{De,Di}$ | 0.95 | 0.95 | 0.95 | 0.95 |
| $s_{Di,De}^{Cy}$ | 2 | 0.02 | 2 | 0.02 |
| $b_{CD}$ | 60 | 70 | 70 | 100 |
| $d_{CD}$ | 0.4 | 0.4 | 0.4 | 0.4 |
| $k_{Di}$ | 200 | 250 | 100 | 120 |
| h | 1.2 | 0.6 | 1.5 | 1 |
| l | 3 | 3 | 3 | 3 |
| $d_{Cy}$ | 1 | 1 | 1 | 1 |
| $\alpha$ | 4 | 4 | 4 | 4 |

### Natural mutation and variation of parameters

In the simulation we introduce a natural mutation to investigate the spontaneous occurrence of Pmel^KO^ mutants due to mutation from the wild-type population (in contrast to artificially introducing them into the tumor). We choose the mutation probability such that the occurrence of a mutant is likely but fixation is not ensured. With m=10e-7, the probability of at least one mutant occurring before the therapy starts at day 14 is approximately 80% but the probability of more than three mutants occurring is only 10% (the growth of the wild-type population is roughly exponential and the occurrence of mutants is a Poisson point process for which probabilities can be calculated).

In Figures 3E, 5C, and S4 the birth rate b_KO_ and natural death rate d_KO_ of the Pmel^KO^ mutants are varied between [0.3,0.55] and [0.01,0.26] respectively to obtain individual fitness values r_KO_=b_KO_-d_KO_ in [0.2,0.45].

## DATA AND SOFTWARE AVAILABILITY

The stochastic-deterministic hybrid algorithm is implemented in C++. The source code can be made available upon inquiry.

