## Supplemental Figures 1-4 and Legends for "Experimental and stochastic models of melanoma T-cell therapy define impact of subclone fitness on selection of antigen loss variants"

**Supplemental figure titles and legends**

**Supplemental Figure S1 (Related to Main Figure 1). Dynamic changes of Pmel protein level in response to METi and therapeutic efficacy of ACT$^{METi}$ against HCmel12$^{METi\text{-}R}$ melanomas.**

**(A)** Experimental outline of generation of METi-resistant HCmel12 variants (HCmel12$^{METi\text{-}R}$). **(B)** Representative images of crystal violet stained culture dishes stained after 6 days of culturing. Cell lines and treatments are indicated. **(C)** Quantification of experiments described in B performed in biological multiplicates (n=5). **(D)** Immunoblots for signaling components downstream of c-Met and β-Actin in indicated melanoma cell lines exposed to vehicle (DMSO) or METi (capmatinib). **(E)** Immunoblots for Pmel and β-actin in indicated melanoma cell lines exposed to vehicle (DMSO), METi (capmatinib) and MEKi (trametinib). **(F)** Flow cytometric analyses of Ngfr (melanoma dedifferentiation maker) surface expression in HCmel12 WT and B16F1 melanoma cells exposed TNF-α or left untreated in combination with vehicle (DMSO) or METi (capmatinib) treatment. **(G)** Experimental outline comparing the therapeutic efficacy of ACT and ACTMETi in mice bearing HCmel12$^{METi\text{-}R}$-#1 melanomas. **(H)** Kaplan-Meier survival curves for the experiments described in G. Log-rank test. *p<0.05. **(I)** Individual tumor growth curves of the experiments described in G. **(J)** Flow cytometry-based quantification of the percentage of Pmel-1 T- cells among peripheral blood leukocytes (PBL) from mice treated as indicated (1 week after onset of ACT regimens). Experiments as described in G. (n=8 for ACT; n=12 for ACT$^{METi}$; unpaired two-tailed t-test ***p<0.001)

**Supplemental Figure S2 (Related to Main Figure 2). Basic characterization of three HCmel12 Pmel$^{KO}$ single cell clones (subclones)**

**(A)** Results from amplicon NGS analyses showing respective frameshift indels in the *Pmel* gene of indicated HCmel12$^{PmelKO}$ single cell clones. **(B)** Immunoblots for Pmel, tyrosinase and β-actin in HCmel12 WT and Pmel$^{KO}$ single cell clones treated as indicated. (C) Experimental outline and tumor growth curves of HCmel12 WT versus mixed HCmel12 Pmel$^{KO}$ single cell clone melanomas in untreated mice.

**Supplemental Figure S3 (Related to Main Figure 3). Optimization of the parameter α.**

(A) Percentage of Pmel$^{KO}$ cells in simulations, taken at tumor size of 10 mm, varying parameter α and initial size of the tumor. Initial portion of Pmel$^{KO}$ cells always at 17.1%. (B) Percentage of Pmel$^{KO}$ cells in simulations, taken at varying tumor sizes and parameters α. Initial tumor of medium size with 17.1% Pmel$^{KO}$ cells.

**Supplemental Figure S4 (Related to Main Figure 5)**

Prediction of enrichment of $Pmel^{KO}$ cells for subcritical initial wild-type tumor and natural mutation at rate m=10e-7 under varying subclone fitness rKO=bKO-dKO and different tumor sizes at time point of harvesting.

Glodde, Kraut et al. Figure S1

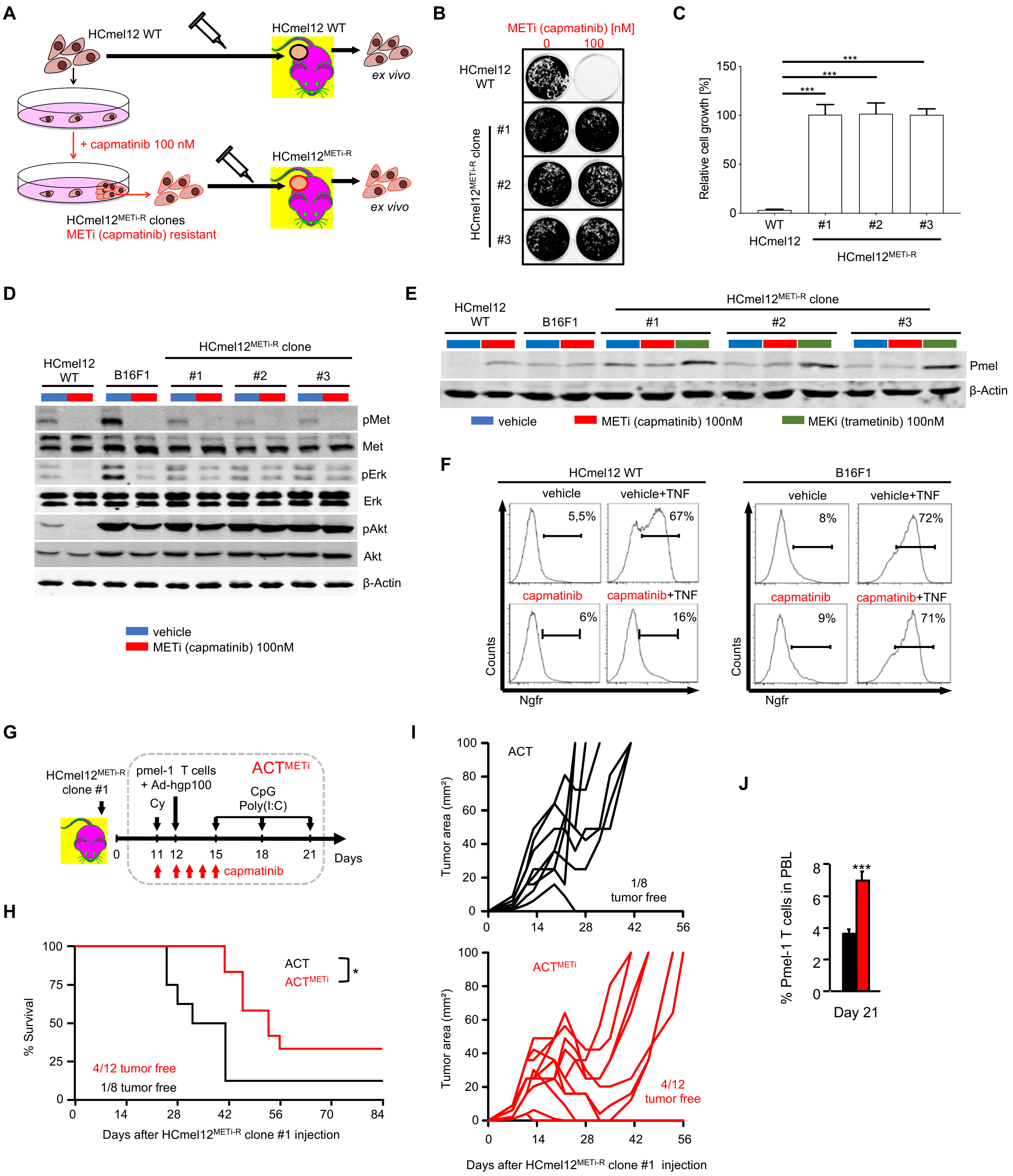

Glodde, Kraut et al. Figure S2

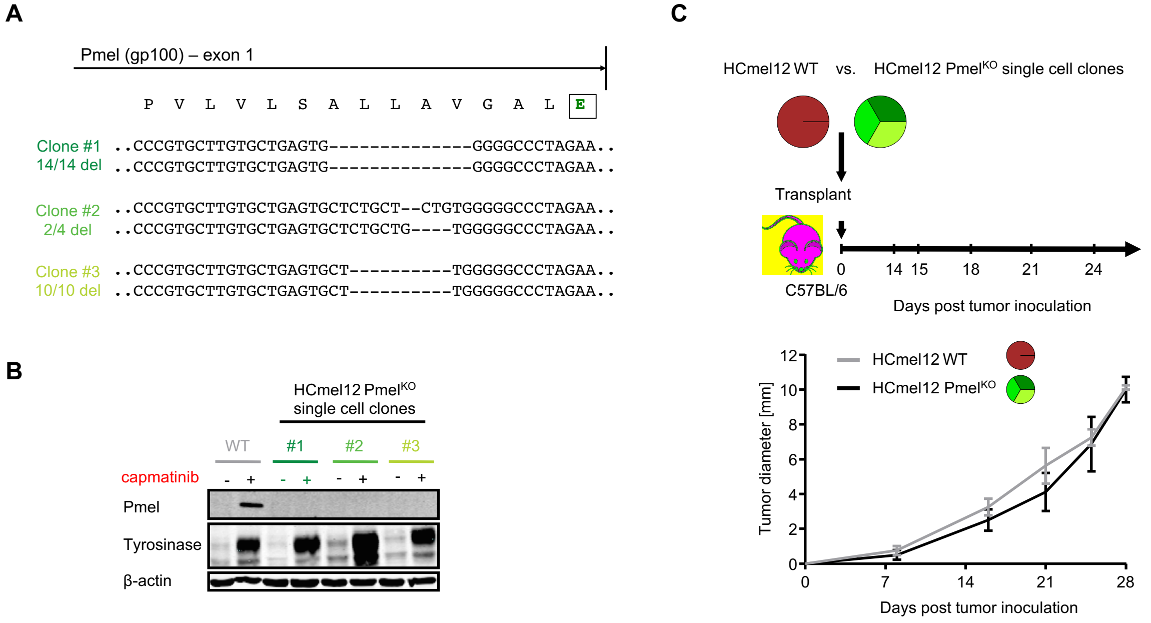

Glodde, Kraut et al. Figure S3

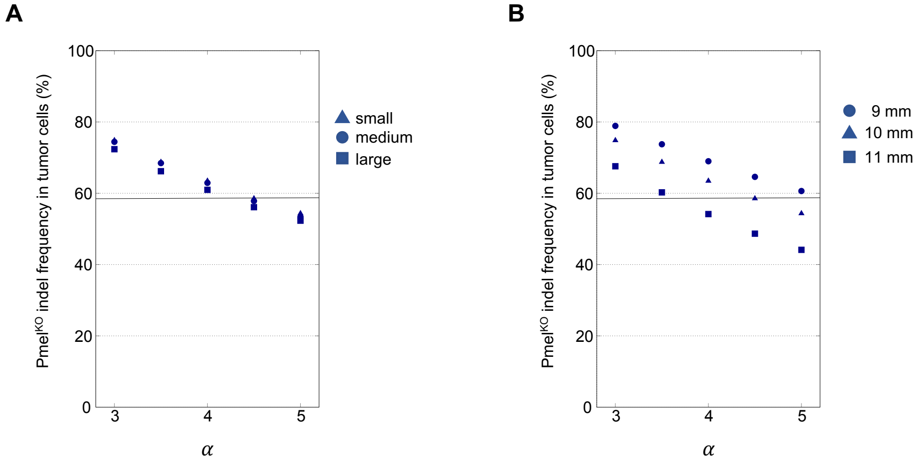

Glodde, Kraut et al. Figure S4

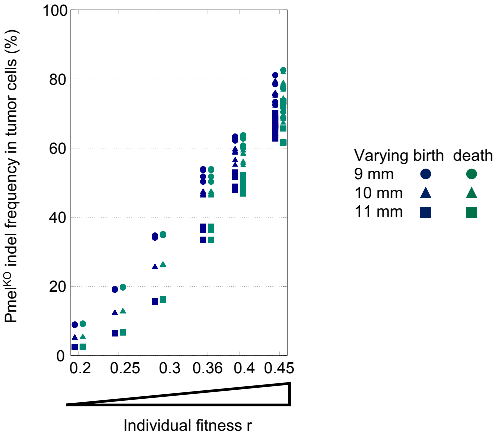
